# Comparative profiling of cellular gait on adhesive micropatterns defines statistical patterns of activity that underlie native and cancerous cell dynamics

**DOI:** 10.1101/2023.10.27.564389

**Authors:** John C. Ahn, Scott M. Coyle

**Affiliations:** Department of Biochemistry, University of Wisconsin-Madison, Madison, Wisconsin 53706, USA; Integrated Program in Biochemistry Graduate Program, University of Wisconsin-Madison, Madison, Wisconsin 53706, USA

## Abstract

Cell dynamics are powered by patterns of activity, but it is not straightforward to quantify these patterns or compare them across different environmental conditions or cell-types. Here we digitize the long-term shape fluctuations of metazoan cells grown on micropatterned fibronectin islands to define and extract statistical features of cell dynamics without the need for genetic modification or fluorescence imaging. These shape fluctuations generate single-cell morphological signals that can be decomposed into two major components: a continuous, slow-timescale meandering of morphology about an average steady-state shape; and short-lived “events” of rapid morphology change that sporadically occur throughout the timecourse. By developing statistical metrics for each of these components, we used thousands of hours of single-cell data to quantitatively define how each axis of cell dynamics was impacted by environmental conditions or cell-type. We found the size and spatial complexity of the micropattern island modulated the statistics of morphological events—lifetime, frequency, and orientation—but not its baseline shape fluctuations. Extending this approach to profile a panel of triple negative breast cancer cell-lines, we found that different cell-types could be distinguished from one another along specific and unique statistical axes of their behavior. Our results suggest that micropatterned substrates provide a generalizable method to build statistical profiles of cell dynamics to classify and compare emergent cell behaviors.

## Introduction

Dynamic behaviors of living systems, from the macroscale to the microscale, are powered through patterns of activity. For a walking human, alternating right-left sequences of pedal contact, arm swinging, and joint articulation produce an emergent walking gait that enables efficient locomotion^1, 2^. Similarly, at the single-cell level, patterns of actin polymerization, filament sliding, and other active processes can coordinate and collaborate to manipulate cell shape to orchestrate complex cell behavior such as phagocytosis or motility^3–5^.

At the molecular scale, metazoan cell migration and adhesion are regulated by signaling through *integrin receptors* that physically couple the binding of extracellular matrix ligands from the environment to the internal cytoskeleton^6, 7^. At the cellular scale, multiple integrin contact sites interface with the specific configuration of signaling networks, cytoskeletal structures and other molecular systems present in the cell to influence the migratory and adhesive behaviors that emerge^8^. This complex interplay between sensory inputs and the biological pathways that process them can cause different cells placed in identical environments to behave in distinct ways; and identical cells in different environments to respond with markedly different behaviors^9^.

However, a quantitative or statistical understanding of the patterns of morphological activity—or “gait” — that occur within a given cell type and how those patterns are modulated by the environment are generally unclear. While mathematical and statistical treatments of gait are commonplace tools to explore such patterns in macroscale biology, where they have great utility in quantifying animal behavior and in diagnosing orthopedic and ergonomic pathologies, these powerful methods are less often applied to single cells^10^. One reason for this has been an emphasis on first identifying the specific molecular components and biochemical mechanisms that give rise to motility in these systems. A second reason is that the morphology of many cells, in particular adherent metazoan cells, lacks the stereotyped anatomy that is commonly used to perform such analyses^11^. That is, while the arms, legs, feet and joints of a human provide concrete and trackable points to build up a description of gait, an adherent metazoan cell’s geometry and sub-cellular anatomy can be vague or rapidly reconfigurable **(Fig. 1A)**.

**Figure 1.**
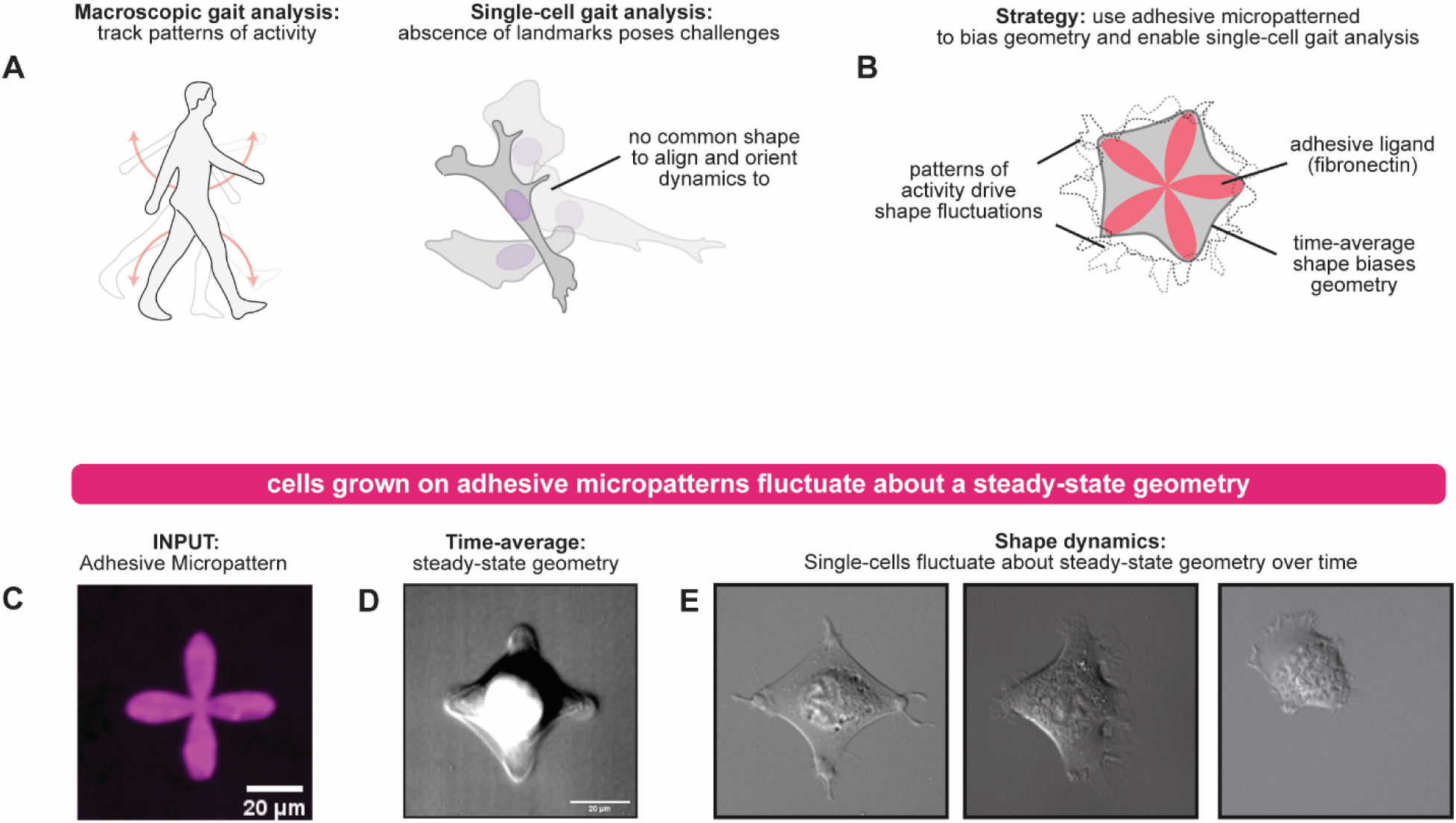
Cells grown on adhesive micropattern islands undergo dynamic shape fluctuations about an average cell shape. (a) Schematic for how gait analysis is used to track patterns of activity in macroscopic biological systems but is difficult to extend to metazoan cells owing to a lack of landmarks and stereotyped geometry. (b) Schematic for strategy of using adhesive micropatterns to bias cell geometry into a configuration suitable for single-cell shape analysis. (c) Fluorescence microscopy image of a fibronectin adhesive micropattern used as a substrate for single-cell gait analysis. The intensity indicates the position on the substrate where the fibronectin ligand has adhered. (d) Time-average image produced from an image stack of a 3T3 cell grown on the adhesive micropattern in (c) over a 12 hours period. (e) Representative images from the time-series used to produce the time-average in (d). The cell geometry is observed to fluctuate substantially from the time-average shape throughout the timecourse.

Indeed, the most successful instances of applying gait analysis to single cells so far have either been done on protozoan cell-types with easily scored sub-cellular anatomy and structure; or cells undergoing such highly stereotyped and reproducible migration that the cell geometry is stable enough for such treatment, such as in migrating fish keratocytes^12–16^. Given the utility of gait analysis for macroscopic biological systems, a general method for performing gait analysis on morphologically plastic metazoan cells would be valuable, as the dynamic behaviors of such single cells provide the foundation of multicellular processes like development, wound healing, and immune responses, and aberrant migration and adhesion contribute to developmental disorders, metastasis, bleeding pathologies, and autoimmunity^17–23^.

Here we develop a general strategy for comparative profiling of cellular gait across different metazoan cell-types and environmental INPUTs. Our strategy is based on quantifying the long-term morphology dynamics of cells grown on different cell-sized micropattern islands of the adhesive ligand fibronectin **(Fig. 1B)**^24^. We find that cells grown on such adhesive micropatterns adopt a steady-state cell geometry in the time-average, but also display shape fluctuations over time about that mean geometry. We develop a computational strategy to digitize these cell morphology dynamics and map them to a polar coordinate (**r,θ**) system to produce morphological signals for quantitative analysis. Inspection of these signals suggests a natural decomposition into short timescale “events”–bursts of highly dynamic activity–and slower timescale meandering in shape variation. We develop specific quantitative and statistical metrics to describe these features and use these to build statistical profiles for specific cell types based on hundreds of thousands of unique measurements in response to a wide range of pattern INPUTs. This allowed us to classify behavior modulation induced by the environment in specific cell lines and identify specific axes of cell behavior that distinguish different cancer cell lines from one another. Because our approach relies only on non-fluorescent imaging of single cells and does not require genetic modification, we anticipate our method may be particularly well-suited to build statistical profiles of cellular gait in rare cell populations or patient-derived samples. Long term, the extension of gait-analysis to metazoan single-cell biology could open up new avenues for clinical diagnostics and guide behavior-informed therapeutic intervention.

## Results

### Cells grown on adhesive micropatterned islands exhibit fluctuations about a steady-state geometry that generates a dynamic morphology signal

Because cells are active systems, they continue to generate dynamics even when biased towards a particular steady-state geometry. To explore the kinds of morphology signals individual cells generate under such conditions, we used DIC microscopy to continuously image individual 3T3 fibroblasts adhered to cell-sized micropatterned islands of fibronectin **(Fig. 1C-E and Fig. S1)**. This approach allowed cell morphology to be easily scored at high temporal resolution (2 frames per minute) without the need for phototoxic fluorescence imaging to be used, enabling timelapse recordings greater than 12 hours to be readily collected for individual cells.

The time average of the image stack for an individual cell’s time series revealed an overall shape resembling the convex hull of the underlying micropattern **(Fig. 1D)**. These time-average shapes resemble the population-average shapes seen when one aligns and averages thousands of different cells grown on separate patterns together^25–27^. However, at any moment in time, a cell was often found to be adopting a geometry that deviated substantially from this time-average shape. For example, while a cell grown on a four-leaf clover island had a diamond-like time average that contacted all four lobes of the pattern, that same cell was observed in shapes with 3-points of contact, shapes that appeared to be polarized between lobes, as well as shapes nearly identical to the time average itself (**Fig. 1E**). Similar phenomena were observed for other micropatterned islands that varied in size or spatial complexity **(Fig. S1)**. These data indicate that while an adhesive micropattern biases the cell to a specific cell geometry, spontaneous cellular activity leads to excursions that drive the cell away from this attractive basin.

### Digitization of morphological signals reveals short and fast timescales underlying cell behavior and defines statistical metrics for analysis

The shape fluctuations we observed for individual cells adhered to micropatterns invite a more detailed and quantitative analysis. Because micropatterned islands are on the order of the size of the cell itself, both the cell geometry and the pattern are well suited to a description in polar coordinates. Thus, we adopted an approach wherein we digitized cell shape for every frame in the recording by calculating the distance **r** from the center of the pattern to the edge of the cell for every angle **θ** between 0 and 360 degrees **(Fig. 2A)**. This maps the time-dynamics of the cell shape from an inconvenient cartesian (**x**, **y**) form into an (**r**,**θ**) “morphological signal”. This (**r**,**θ**) version of the signal is particularly convenient as it is easy to align across individual cells and facilitates the application of many visualization and statistical techniques to interrogate the content of the signal.

**Figure 2.**
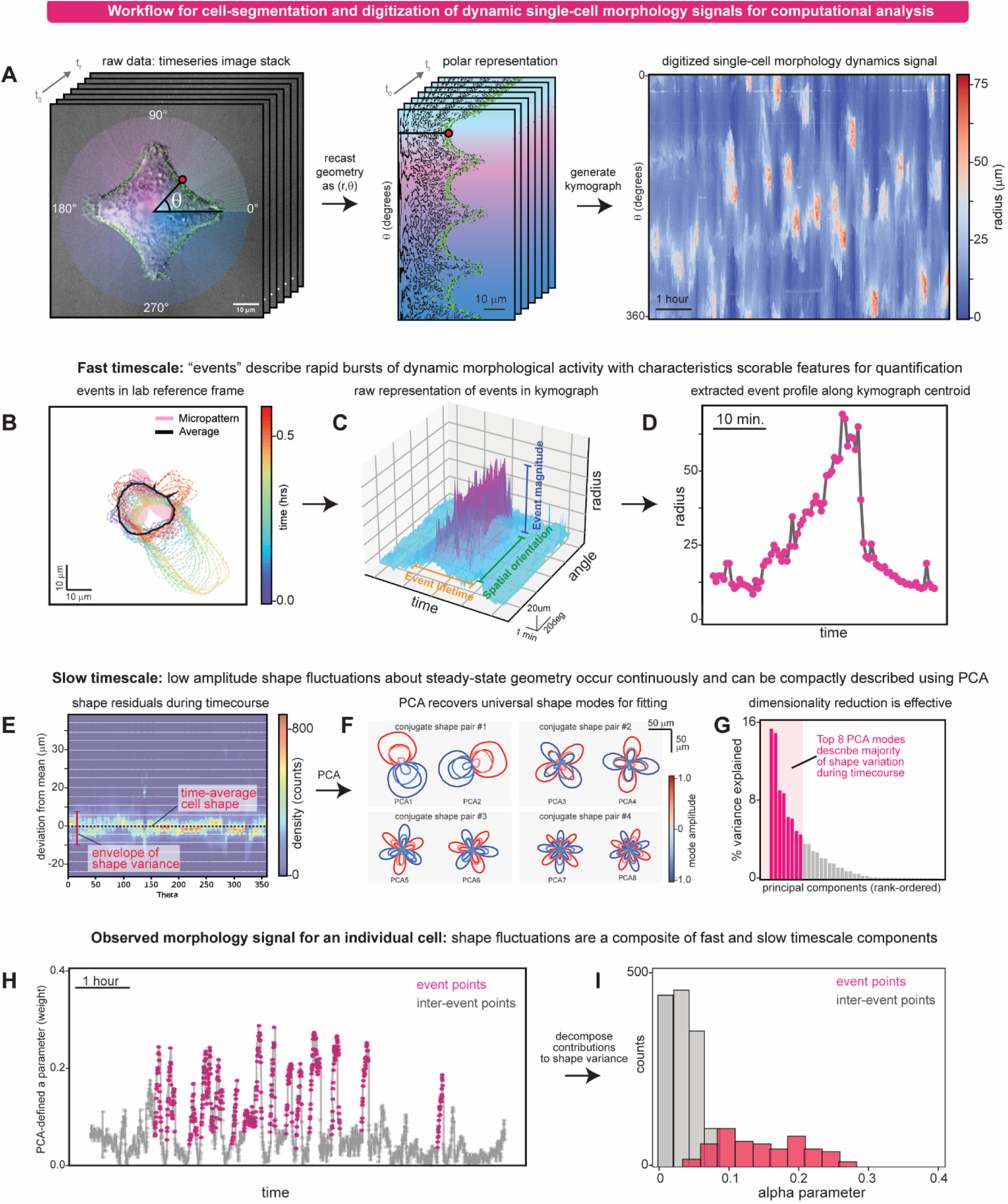
Digitization of single-cell morphology signals defines statistics for fast and slow timescale contributions to shape dynamics. (a) Workflow for digitization of cell morphology signals generated by cells grown on adhesive micropatterns, as applied to a representative cell. Using the center point of the micropattern as an origin, the cell shape is mapped from cartesian coordinates to an r,q description that describes the cell’s radius at each possible angle from 0-360 degrees. By performing this for each image in the time-series, a compact description of the cell’s geometry over time is obtained. The content of these timeseries can be intuitively viewed as a kymograph, which shows bursts of dynamic activity we term ‘events’, within the cell’s morphological signal. (b) Visualization of a single “event” from the kymogrpah shown in (a), showing the contour of the cell over time. For this event, it clearly corresponds to a large extension of the cell geometry followed by retraction. (c) Structure of the event from (b) using the kymograph as a reference point. In the r,q reference frame the events appears as a burst of activity with an easily scored lifetime, amplitude, and orientation. (d) Two-dimensional representation of the profile of the event from (c) by projecting the data along its central angle. (e) Two-dimensional histogram depicting the distribution of shape residuals (observed shape – mean shape) for the cell from (a). Most observations fall within a 10 micron envelope about the cell’s steady state geometry. (f) Real-space depictions of the top 4 conjugate pair shape modes obtained by performing PCA analysis on the data in (e). Colors depict the effect on shape arising by adding (red) or subtracting (blue) the weighted shape mode from the average shape. Note that the PCA-derived shapes correspond to increasingly higher-order Fourier shape modes (1,2,3,4 order depicted). (g) Histogram showing the percentage of shape variance explained by each individual shape mode obtained by PCA decomposition of the data in (e). The top 8 shape modes describe >80% of the observed shape variance in the dataset and were selected to use for weight calculation, fitting, and dimensionality-reduction. (h) 1-dimensional representation of the morphological signal from (a) using only the weight amplitude (denoted as α) for the first conjugate pair of shape modes in (f). Points colored in pink correspond to time points associated with “events” from the kymograph in (a). Note that high amplitude shape mode usage appears to correspond to when events occur. (i) Partitioning of the overall distribution of α values from (h) into “event” and “inter-event” timepoints. Inter-event distributions have low α values compared to a right-shifted distribution for the event distribution.

Close inspection of a representative morphological signal from a 3T3 cell grown on a 30µm wide k2 micropattern suggests several axes to quantify and unpack in our analysis. First, the cell’s morphology signal can be unwrapped and displayed as a kymograph, providing an intuitive visualization of how a cell’s shape fluctuations are distributed in both space and time during the experiment **(Fig. 2A)**. For the representative 3T3 signal, we observe a structure in which discrete “events”–bursts of localized spatiotemporal activity–occur periodically throughout the signal. In between events, the cell’s shape continues to fluctuate but with much lower amplitude and without obvious biases in its spatiotemporal character. This suggested that a cell’s morphology signal can be at least partially decomposed into two different components: 1) a description of the statistical properties of the ‘events’; and 2) a description of the global and inter-event shape fluctuations.

To create metrics for characterizing the “events” within a cell’s morphological signal, we directly mined features of the kymograph itself **(Fig. 2B)**. For any individual event, we defined an associated duration, amplitude, orientation, and width for that event **(Figure 2C-D)**. Moreover, because multiple events are observed within a single morphological signal, individual events and their appearance over time can be used to define an event density (events/hour) and associated average inter-event time as well.

To quantitatively characterize the morphological dynamics between events, we also inspected and analyzed the distribution of variation about the cell’s mean shape. For any observed shape vector, the difference between that shape and the time-average shape defines a residual shape **(Fig S2)**. By aggregating these residuals across all timepoints and inspecting their distribution across θ values, we observed that the bulk of observations generally fell within an envelope of values about the mean shape **(Fig. 2E)**. To determine what sorts of shape vectors best described these fluctuations, we used principal component analysis (PCA) to identify a set of 8 shape modes that could explain >70% of the variance **(Fig. S2)**. Mapping these shapes from (**r**,**θ**) back to the lab reference-frame revealed that the modes we recovered appeared to naturally correspond to conjugate pairs of Fourier shape modes **(Fig. 2F)**. Generally, the lowest frequency modes explained the greatest proportion of the variance, describing simple polarization of the mean shape, with higher frequency modes playing a minor role that allows for more complex geometries **(Fig. 2G)**. This universal collection of shape modes can be used to describe the shape fluctuations about a cell’s steady-state geometry on any pattern or for any cell type **(Fig. S2)**.

Using these two descriptors, we can begin to see how an individual cell’s morphological signal emerges as an interplay between these fast events and slower timescale shape fluctuations. For the representative 3T3 cell, plotting the magnitude of the first conjugate pair of shape modes (here termed **α**) over time provides a 1-dimensional timeseries approximating the cell’s morphology dynamics **(Fig. 2H)**. Large spikes in the magnitude of the **α** parameter coincided with events seen in a cell’s morphology kymograph **(Fig. 2H and Fig. S2)**. This suggested that the discrete events are a large contributor to the overall shape variation observed during the timecourse. Thus, we compared the distribution of **α** values during burst events to the distribution of **α** values occurring outside the events. Indeed, this partitioning revealed a low-magnitude **α** distribution outside of events, and a higher right-shifted magnitude for the **α** distribution within events **(Fig. 2I and Fig. S2)**. As such, the relative contribution of the two **α** distributions to the overall variation provides another useful metric for describing the underlying structure of a cell’s morphological signal.

Having developed a strategy for digitization of a cell’s morphology signal and having defined metrics to classify different discrete (structure of events) and continuous (**α** distributions) elements of its behavior, we will now begin to apply these quantitative tools to ask how a cell’s morphological signal is affected by environmental inputs or underlying cell-type.

### The spatial structure of a micropattern island modulates the statistics underlying a cell’s morphological signal and emergent behavior

The preceding section developed a pipeline for digitizing cell dynamics on micropatterned islands and associating quantitative metrics and statistics with the resulting emergent morphological signal. A natural question that arises is how the morphological signal that we observe is affected by the underlying structure of the micropattern island itself. Because micropatterning allows manipulation of the spatial structure of the fibronectin INPUT to the cell at micron length-scales, it is straightforward to construct patterns that emulate different environmental features a cell might experience naturally, such as the overall availability of ligand or the number of polarization directions available. Thus, we mathematically generated a family of micropatterned fibronectin island INPUTs using the polar equation for the harmonics of a circle (*r = d + a•cos(kθ)*), which systematically allowed us to sample different size (radius parameters, defined by *a* and *d*) or spatial complexities (*k*, number of petals). This family of patterns was well suited to our analysis as the structure of the pattern is defined in the same polar coordinate system that we use to describe the cell shape itself, allowing us to look for correlations between pattern structure and cell behavior **(Fig. 3A)**.

**Figure 3.**
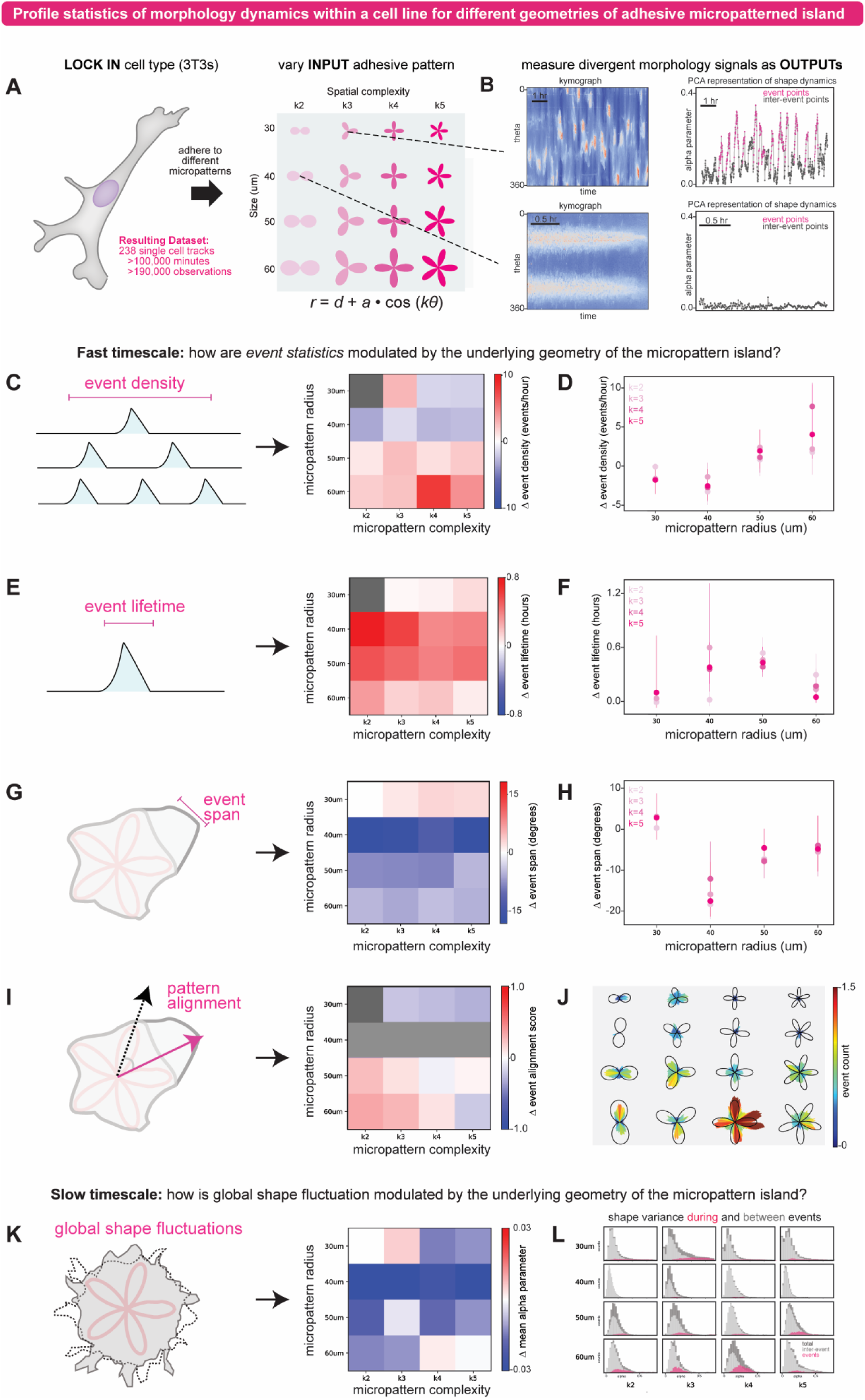
The spatial structure of a micropattern island modulates the statistical features of a cell’s dynamic morphology signal. (a) Schema depicting experimental design for interrogating effects of micropattern geometry on 3T3 morphology signals. 3T3 cells were imaged on different micropatterns defined by the harmonics of a circle, such that the spatial complexity and radius of the pattern were independently controlled. The resulting single-cell tracks were digitized for analysis using the workflow from Fig. 2 and aggregated by INPUT pattern for statistical comparison between different conditions. (b) Examples of divergent morphological signals (presented both as kymographs and α time courses) observed when 3T3 cells were grown on different micropattern substrates. Top: a highly dynamic morphological signal observed from a cell grown on a 30um k3 pattern. Bottom: a cell with very little dynamic content from a cell grown on a 40 um k2 pattern. (c) Relative “event density” statistics across the panel of INPUT patterns, using the 30 um k2 pattern as the reference for comparison by the “shared control bootstrapping” method. (d) Plot of the data in (c) showing the trends observed when micropattern radius is varied for each level of spatial complexity (k2, k3, k4, k5). Values and error bars derived using the “shared control bootstrapping” method (see also Fig. S3). (e) Relative “event lifetime” statistics across the panel of INPUT patterns, using the 30 um k2 pattern as the reference for comparison by the “shared control bootstrapping” method. (f) Plot of the data in (e) showing the trends observed when micropattern radius is varied for each level of spatial complexity (k2, k3, k4, k5). Values and error bars derived using the “shared control bootstrapping” method (see also Fig. S3). (g) Relative “event span” statistics across the panel of INPUT patterns, using the 30 um k2 pattern as the reference for comparison by the “shared control bootstrapping” method. (h) Plot of the data in (g) showing the trends observed when micropattern radius is varied for each level of spatial complexity (k2, k3, k4, k5). Values and error bars derived using the “shared control bootstrapping” method (see also Fig. S3). (i) Relative “pattern alignment” statistics for an event across the panel of INPUT patterns, using the 30 um k2 pattern as the reference for comparison by the “shared control bootstrapping” method. (j) Plot of the data used to generate (i) projected onto the underlying shape of the micropattern for direct visualization of event-to-pattern alignment. (k) “Global shape fluctuation” statistics (from α distributions) across the panel of INPUT patterns, using the 30 um k2 pattern as the reference for comparison by the “shared control bootstrapping” method. (l) Partitioning of the overall distribution of α values from (k) into “event” and “inter-event” timepoints across the panel of different INPUT patterns.

We prepared adhesive fibronectin micropatterns based on a matrix of 4 different radii (30,40,50,60 μm) and 4 different spatial complexities (k-values 2, 3, 4 and 5), performed extended timelapse imaging of 3T3 cells grown on each pattern, and digitized their morphological signals for analysis. Our 3T3 dataset contained a total of 238 cells, spanning 107,382 minutes, for a total number of 196,949 unique observations across the matrix. Because we acquired signals for multiple cells grown on each micropatterned INPUT, the statistical features of their resulting signals could be pooled to generate high credence descriptors of the observed behaviors.

Using the pipeline developed in the previous section, we first asked how the nature of fast-timescale morphological “events” were affected by the geometry of the pattern INPUT. We immediately noticed that the overall event density (number of events/hour) differed dramatically across the panel of INPUTs. For example, cells grown on 40 micron patterns had a very low event density–less than 1 event per hour in some cases; while in contrast, cells grown on larger, more-spatially complex patterns (e.g. 60 um k4 and k5) had much larger event densities, greater than 10 per hour in some cases **(Fig. 3B-C and Fig. S3)**. Thus, the geometry of the adhesive micropattern appeared to be giving rise to markedly different behaviors in the 3T3 cells.

To unpack these observations further, we more closely inspected how specific statistical features of these morphological events changed as a function of size and spatial complexity. To help compare the changes we observed, we used the statistics we collected from cells grown on 30 um k2 patterns as a reference point. The associated raw data and raw metrics, which provide a complementary view into this analysis, are included in Fig. S3. With respect to “event density”, we observed an initial small dip in density as cells moved from 30 um to 40 um patterns (k2,3,4,5 Welch p-values = <0.01, 0.20, 0.01, 0.03), before increasing as the radius increased to 50 (k2,3,4,5 Welch p-values = 0.39, 0.04, 0.25, 0.13) and then 60 um (k2,3,4,5 Welch p-values = 0.16, 0.32, <0.01, 0.11) **(Fig. 3C-D)**. This trend held regardless of the underlying spatial complexity (k-value) of the pattern, although the increase was more pronounced for patterns with high k-value compared to those with lower k-value. This check-mark shaped response to pattern radius we observed appears to qualitatively agree with the extent to which 3T3 cells were able to fully spread across the micropattern: on 30 micron patterns most cells were incompletely spread; on 40 micron patterns most cells were fully spread with little remaining micropattern available to adhere to; and on 50- and 60-micron patterns, cells were fully spread but with additional adhesive surface available to explore or sample.

In addition to event density, we observed that the nature of the pattern (radius and spatial complexity) modulated statistical features of individual events themselves. With respect to radius, the average lifetime of events was longest for 50 micron patterns (k2,3,4,5 Welch p-values = <0.01, <0.01, <0.01, <0.01; compared to the 30 um k2 distribution), followed by 40 micron (k2,3,4,5 Welch p-values = 0.32, <0.01, 0.08, <0.01; compared to the 30 um k2 distribution), 60 (Welch p-values = <0.01, 0.09, <0.01, 0.10; compared to the 30 um k2 distribution) micron and finally the shortest-lived events for 30 microns (k2,3,4,5 Welch p-values = n.a., 0.55, 0.37, 0.50 compared to 30 um k2) **(Fig. 3E-F)**. The span parameter of these events—how wide the morphological extension was—varied substantially across the different patterns, with 40 micron patterns inducing significantly narrower event excursions (k2,3,4,5 Welch p-values = <0.01, <0.01, <0.01, <0.01; compared to the 30 um k2 distribution) compared to 30,50, or 60 micron patterns **(Fig. 3G-H)**. These results mirror our earlier observation that 3T3 cells are more event-dense when incompletely spread (30 micron) or have excess adhesive ligand available (50 and 60 microns). Effects of a pattern’s spatial complexity on these event statistics were less obvious, but still detectable in some cases. For example, higher k-value patterns (more petals) were generally associated with shorter lived events. However other parameters like the “span” of the pattern were not obviously influenced by the k-parameter.

Given these observations, we wondered whether the spatial structure of the pattern might have stronger influence on the spatial orientation of events as opposed to their general features. To explore this, we projected the orientation of an event (defined as the θ values associated with the span of an event) onto the underlying structure of the micropattern to create a histogram for visualizing event alignment **(Fig. 3I-J)**. Inspection of these histograms qualitatively suggested that, for cells with high event densities, the events were more spatially aligned with the micropattern as the size of the pattern increased and the spatial complexity decreased.

To analyze this quantitatively, we computed an alignment score based on event centroid angular overlap with the underlying micropattern and binned the scores for each pattern together **(Fig. 3I-J)**. Because the 40 micron patterns induced very few events, their alignment scores were too low confidence and were excluded from detailed analysis. Nonetheless, a clear trend towards better event/pattern alignment was observed for increasing pattern radius (k2 at 50µm, k3 at 50µm, k2 at 60µm, k3 at 60µm Welch p-values = <0.01, <0.01, 0.05, 0.09; compared to the 30 um k2 distribution). Moreover, a decrease in alignment score was observed as the spatial complexity of the pattern increased. An intuitive hypothesis for the interpretation of these data is that a cell’s ability to effectively orient along a micropattern reflects a sort of spatial resolution inherent to the cell. That is, events become better oriented if an open area of adhesive ligands is large enough to polarize along, and the combination of spatial complexity and pattern radius determines whether or not that area requirement is being satisfied. By probing both radius and spatial complexity independently, our micropatterning assay was able to identify this inflection point for the 3T3 cells, which could be different depending on cell-type or cell physiology.

Finally, we examined how the statistics describing cell behavior *between* events—that is slow-timescale meandering—was affected by the micropattern. Using our universal collection of PCA shape modes **(Fig. 2F and Fig. S2)**, we fit our observations and decomposed the global α parameter distribution associated with each micropattern INPUT into its associated event and non-event distributions **(Fig. 3K-3L)**. We found that although the event α distribution differed considerably across different pattern INPUTs—which makes sense given the differences in event statistics we described above—we observed only very minor differences to the inter-event α distribution. That is, for the 3T3 cells we inspected, the statistics describing the slow timescale meandering behavior between events were generally similar regardless of the pattern tested.

Taking together, our query of 3T3 behavior across a range of different micropatterned INPUTs suggests that the structure of the micropattern INPUT largely modulates the event, but not inter-event, statistics underlying the cell’s emergent morphology signal. Properties such as event lifetime, frequency, span, and orientation are affected by the nature of the pattern, with size (pattern radius) appearing to play a larger role than spatial complexity (petal number). In contrast, the overall morphological fluctuations and shape meandering occurring between these events was not strongly affected. Thus, using adhesive micropatterns, we can build a statistical profile that captures a given cell-type’s response to a range of different environmental INPUTs.

### Profiling cancer-cell lines on micropatterned islands identifies different statistical axes associated with divergent morphological signals and cell dynamics

The preceding section demonstrated that a single cell-type can display a complex set of morphological responses to a defined set of structured micropattern INPUTs, and that these differences can be used to build a statistical profile for how pattern INPUTs are mapped into morphological signal OUTPUTs. Given this, we next wondered whether our assay could be used to profile different cell-types for comparison, and whether we could identify specific statistical axes defining the key differences underlying divergent behaviors between cell-types. To this end, we selected four cell lines—Hs578t, BT549, MDA-MB-436, and MDA-MB-231—from an established panel of triple negative breast cancer cell lines with mesenchymal or luminal morphology for investigation^28^. Some of these lines, such as MDA-MB-231, have been previously noted for their high levels of aberrant motility and are frequently termed “hyper-metastatic” in the literature^29^.

We performed extended timelapse imaging for each cell line grown on adhesive fibronectin micropatterns with a radius of 50 microns and 4 different k values (petal numbers 2, 3, 4 and 5) **(Fig. 4A)**. This subset showed a good range of statistical diversity in the 3T3 experiments, making it a natural starting point for comparison across different cell lines. Single-cell morphology signals were digitized as before, and signals from pattern-matched cells and cell-lines were binned for statistical analysis **(Fig. 4B)**. In parallel we analyzed 3T3 cells grown on identical conditions to use as a non-cancerous “out-group” reference cell-line for comparison. Our cell panel dataset contained a total of 396 cells, spanning >218,000 minutes, for a total number of >247,000 unique observations across the matrix. The associated raw data and statistics have been included in Fig. S4. Different cancer-cell lines from the panel exhibited obvious qualitative differences in behavior **(Fig. 4B)**. For example, the morphology signal kymographs for MDA-MB-231 had high levels of fast timescale activity, while MDA-MB-436s often exhibited extended periods of slow activity.

**Figure 4.**
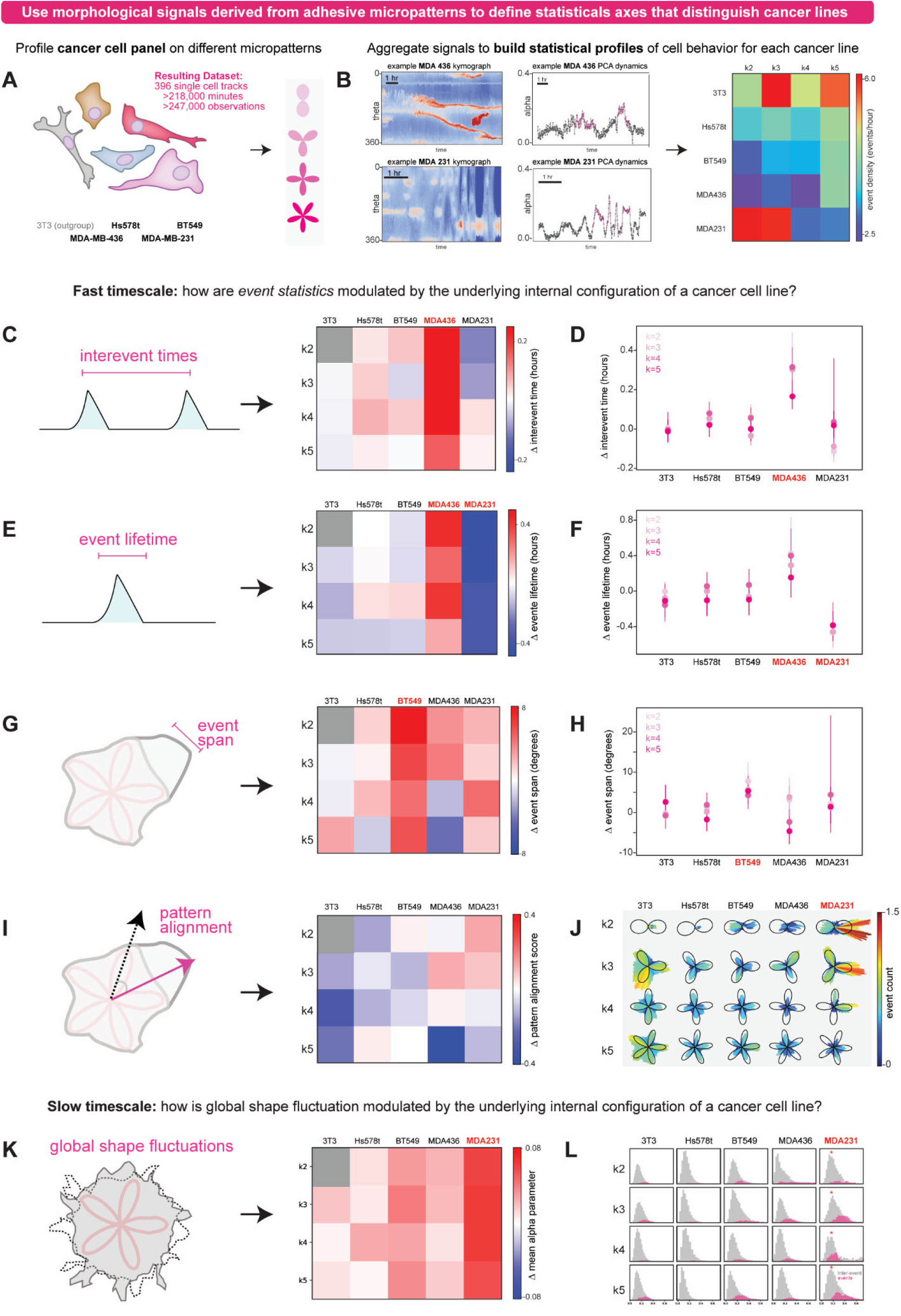
Profiling cancer-cell lines on micropatterned islands identifies different statistical axes associated with divergent morphological signals and cell dynamics. (a) Schema depicting experimental design for building statistical profiles for different cancer cell lines based on the morphology signals they generate when adhered to different micropattern geometries. Cell lines were imaged on 50 um micropatterns across 4 levels of spatial complexity (k2, k3, k4, k5). The resulting single-cell tracks were digitized for analysis using the workflow from Fig. 2 and aggregated by cell-type and INPUT pattern for statistical comparison between different conditions. (b) Examples of divergent morphological signals (presented both as kymographs and α time courses) observed when different cancer cells are grown on different micropattern substrates. Top: an example morphological signal observed from a MDA438 cell Bottom: an example morphological signal from an MDA231 cell. (c) Relative “interevent time” statistics across the panel of cell lines and input patterns, using the k2 3T3 data as an outgroup reference for comparison by the “shared control bootstrapping” method. (d) Plot of the data in (c) showing the trends observed when micropattern spatial complexity is varied for each cell line tested. Values and error bars derived using the “shared control bootstrapping” method (see also Fig. S4). (e) Relative “event lifetime” statistics across the panel of cell lines, using the k2 3T3 data as an outgroup reference for comparison by the “shared control bootstrapping” method. (f) Plot of the data in (e) showing the trends observed when micropattern spatial complexity is varied for each cell line. Values and error bars derived using the “shared control bootstrapping” method (see also Fig. S3). (g) Relative “pattern alignment” statistics across the panel of cell lines, using the 3t3 k2 pattern as the reference for comparison by the “shared control bootstrapping” method. (h) Plot of the data in (g) showing the trends observed when micropattern spatial complexity is varied for each cell line. Values and error bars derived using the “shared control bootstrapping” method (see also Fig. S3). (i) Relative “pattern alignment” statistics for an event across the panel of cell lines, using the 3t3 k2 pattern data as the reference for comparison by the “shared control bootstrapping” method. (j) Plot of the data used to generate (i) projected onto the underlying shape of the micropattern for direct visualization of event-to-pattern alignment across the different cell lines tested. (k) “Global shape fluctuation” statistics (from α distributions) across the panel of INPUT patterns, using the 3t3 k2 pattern data as the reference for comparison by the “shared control bootstrapping” method. (l) Partitioning of the overall distribution of α values from (k) into “event” and “inter-event” timepoints across the panel of different cell lines.

Similar to our earlier analysis of the 3T3s, we used our quantitative statistical metrics for describing events and global shape variation to unpack the differences in morphological behavior for the different cancer-cell lines across specific parameters. We found that although all of these cell lines were associated with metastatic breast cancer, they could be distinguished from one another on the basis of specific statistical metrics that were uniquely affected within each line. For some cell lines, these differences manifested specifically in the nature of the statistics of fast timescale events. For example, MDA-MD-436 cells had unusually long events (k2,3,4,5 Welch p-values = 0.021, <0.01, <0.01, 0.186; compared to 3T3 30 um k2 distribution) and high inter-event waiting times (k2,3,4,5 Welch p-values = <0.01, <0.01, <0.01, <0.01; compared to 3T3 30 um k2 distribution), meaning that events occurred less frequently but each event was markedly longer in duration **(Fig. 4C-F)**. In another example, we found that while some cell-lines produced larger “span” events for certain k-value patterns, BT549 cell lines were unique in that they produced events with significantly larger “span” than the other cell lines across all k-values tested (k2,3,4,5 Welch p-values = <0.01, <0.01, <0.01, <0.01; compared to 3T3 30 um k2 distribution) **(Fig. 4G-H)**. This means that the nature of the BT549 events makes them statistically wider and bulkier on average than any of the other cell-lines we examined.

In other cases, global and inter-event statistical descriptors, like the α parameter distribution, could capture differences between cell lines. For example, MDA-MB-231s had a significantly higher average non-event α parameter than any of the other cell lines we examined (k2,3,4,5 Welch p-values = <0.01, <0.01, <0.01, <0.01; compared to 3T3 30 um k2 distribution) **(Fig. 4K-L)**. This means that MDA-MB-231s have a greater baseline shape variation than the other cell-lines tested, even when no events are occurring. Indeed, while MDA-MB-231s often had fewer and shorter events than other cell-lines (k2,3,4,5 Welch p-values = <0.01, <0.01, <0.01, <0.01; compared to 3T3 30 um k2 distribution), they nonetheless had the highest overall shape variation of any cell line we inspected. Interestingly, when MDA-231s did perform events, they generally polarized the cell strongly along the orientation of the micropattern structure (k2,3,4,5 Welch p-values = 0.02, 0.04, 0.813, 0.05; compared to 3T3 30 um k2 distribution) **(Fig. 4J)**. Thus, the statistical profile of MDA-MB-231s that emerges is one in which a high level of steady-state shape fluctuation is punctuated by highly polarized events. These observations are consistent with the behavior of MDA-231s grown on narrowly-connected two-island patterns^24^, where they exhibit large shape fluctuations and high rates of translocation between the two islands.

Taken together, our results indicate that the morphology signals emitted by different cell-lines grown across different micropatterned INPUTS can generate statistical profiles that can be used to describe and distinguish the behaviors of different cell types. Although all the cells tested were derived from malignant triple-negative breast cancers, the statistics underlying their dynamic morphologies could nonetheless be markedly different. These differences manifested not only as changes in magnitude of some common statistics, but also in terms of which parameters were specifically affected, and even in terms of which timescales (fast events versus slow meandering) were affected. We hypothesize that the underlying internal configuration inherent to each cell-type we tested can manifest as changes to a specific statistical axis underlying a cell’s morphology signal, and careful analysis of these signals can identify the specific axes that are changed.

## Discussion

Here, we have used cell-sized adhesive micropatterns as a platform for building statistical profiles describing the patterns of activity underlying a cell’s emergent dynamics. We found that by biasing the cell’s steady-state shape to a specific geometry, micropatterns provided a straightforward way to digitize the shape fluctuations about this steady-state shape for statistical analysis. By mapping these dynamics onto a polar coordinate description, we could treat a cell’s morphology signal as a kymograph in the (**r**,**θ**) reference frame and analyze its content in this space. This revealed a natural partitioning of the dynamics into short-timescale discrete “events” and a slower timescale meandering of shape between these events. We developed a set of metrics to describe the statistical features of these fast morphology events (event frequency, lifetime, span, alignment) and inter-event shape fluctuations (PCA-derived alpha distributions).

We used these metrics to ask how morphological signal was affected by the geometrical features of the pattern INPUT, using 3T3s as a test cell line. This revealed that 3T3 *event* statistics were strongly modulated by different INPUT patterns, but that the baseline shape fluctuations and global meandering were generally unaffected. We then extended this approach to build statistical profiles for members of a panel of triple-negative breast cancer cell lines. This revealed that different metastatic breast cancer cell lines could exhibit aberrant dynamics with markedly different statistical underpinnings. For example, MDA-MB-231s showed large shifts to its global shape variation parameters, whereas MDA-MB-436s were largely defined by changes to the event structure, producing much longer and more infrequent events. Thus, our method provides a means by which the statistical features of single-cell behavior can be efficiently digitized, aggregated, and compared for analysis.

The approach we take here treats the emergent behavior of a cell–the dynamics to its shape envelope–as an OUTPUT arising from an extraordinarily complex set of internal molecular processes that can react to specific features of the pattern INPUT. By choosing metrics which are based on DIC microscopy and not rooted in fluorescence-based signals or detailed molecular markers, our approach avoids phototoxicity and can yield timeseries data constrained only by interference from cell-division events. In this way, our approach may be especially useful for building statistical profiles of cell-lines when only a small number of cells are available, or genetic modification is infeasible or unwanted; for example, in patient-derived samples or rare populations of primary cells.

At the same time, the lack of molecular detail explaining the biochemical origin for the statistical phenotypes we observe at present limits the scope of our interpretation of these data. For example, what is the molecular basis for longer event lifetimes in some cells versus changes to global shape fluctuation seen in others? While many specific hypotheses could be possible, such as RhoGEF activity^30, 31^, integrin expression level and repertoire^32, 33^, or metabolite availability and usage^34, 35^, the number of genomic differences between the different cell lines we inspected is too great to pinpoint any specific cause at this stage^36, 37^. However, the approach we have developed here will be well-suited to future studies aimed at identifying specific molecular drivers to each statistical axis we identify. For example, a CRISPRi/a screen^38, 39^ in an otherwise isogenic cell-line could be used to identify genes whose dysregulation is associated with higher rates of shape fluctuation, longer event lifetimes, or greater event frequencies.

Long-term, we anticipate this approach could be used to map statistical profiles built based only on morphology dynamics to an associated set of genetic legions that are commonly associated with that aberrant phenotype. Such knowledge could help pinpoint specific causal drivers of metastasis operating in patient-derived cell-lines that are often not immediately obvious from a catalog of genome mutations^40–42^. As such, a cell’s own dynamic behavior may, through its statistics, have the potential to report out the anomalies and pathologies lurking within.

## Acknowledgements.

We thank members of the Coyle Lab and Weeks Lab for advice, helpful discussions, and critical reading of the manuscript.

## Author contributions

JA and SMC conceived the overall project. The experimental plan was implemented by JA, and SMC. JA collected all the data in the paper. JA and SMC prepared the figures and wrote the manuscript. SMC supervised all aspects of the work.

## Methods

### Micropatterning

We employed the light-induced molecular adsorption (LIMAP) method for micropatterning ^43^. 35mm Glass bottom petri dishes (MatTek P35G-1.5-20-C) were exposed with oxygen plasma in preparation for passivation. For adsorption of the anti-fouling coating agent, 0.1 mg ml^-1^ PLL(20)-g[3.5]-PEG(2) (SuSoS CHF9,600.00) solution was added for 1 hour. The dish was washed five times with Milli-Q purified water. A 1:5 ratio of PLPP photoinitator gel to 70% ethanol (Alveole) was added to the microwell and dried at room temperature for 1 hour. To create the micropatterns, the well was exposed to UV light at a dosage of 30mJ mm^-2^, and excess gel was washed away with 5 Milli-Q purified water washes and 5 Dulbecco’s Phosphate Buffered Saline (DPBS) solution (VWR L0119-0500) washes, with the last volume of DPBS left to incubate for 5 minutes to rehydrate the substrate. The wells were then incubated with 10µg ml^-1^ fibronectin (Sigma-Aldrich F1141-5MG) and 10µg ml^-1^ NeutrAvidin (Invitrogen 84607) for five minutes and then washed 5 times with DPBS. PEGs were added at 0.1 mg ml^-1^ for another 1 hour incubation before being finally washed with water for 5 times.

### Cell culture

3T3 mouse fibroblast cells (ATCC CRL-1658) were cultured in Dulbecco’s Modified Eagle’s Medium (DMEM) (Sigma-Aldrich D6429) with 10% fortified calf bovine serum (Cytiva Life Sciences SH30396.03) and 1% penicillin-streptomycin (ThermoFisher 15140122). Cells were grown in a 5% CO_2_ incubator at 37°C up to 90% confluence before being washed and passaged. Adherent cells were washed with PBS at each passage and detached from the flask surface by incubating with TrypLE (ThermoFisher Scientific 12604021) for 5-10 minutes at 37°C. TrypLE was quenched with fresh DMEM media, and cells were resuspended and plated into new flasks with fresh DMEM. The following breast cancer cell lines were all cultured in DMEM with 10% fetal bovine serum (Fisher Scientific SH30396.03) and 1% penicillin-streptomycin in a 5% CO_2_ atmosphere at 37°C: MDA-MB-231 (ATCC HTB-26), MDA-MB-436 (ATCC HTB-130), Hs 578T (ATCC HTB-126), and BT-549 (HTB-122).

### Microscopy

Measurements were obtained in time-lapse mode for up to 16 hours on a Nikon Eclipse Ti2 microscope. Throughout the measurements, samples were incubated in a Tokai Stage Top Incubator that maintained a temperature of 37°C and an atmosphere of 5% CO_2_. Bright-field images were acquired every 30, 40, or 60 seconds.

### Image analysis

Custom python scripts for specific image-analysis workflows are outlined below.

#### Encoding polar descriptions of cell outlines

Binary masks of the cells were calculated using a canny filter from python OpenCV packages. The center point of each cell was identified, from which a 360° radial sweep was conducted, resulting in a 2D array of pixel intensities for each surveyed degree along the length of the pixel survey radius. From each surveyed 1D array of pixel intensities, the radius of the cell outline was calculated based on a 95% threshold in which 95% of all the pixels with maximum intensities were located.

#### Kymograph analysis

2D arrays of cell radii along theta values were collected for each point of time for each cell over the length of their microscopy time course. Heatmaps of these arrays were generated using the matplotlib.pyplot package, from which contours were identified using OpenCV packages. These contours were identified as morphological events and used to calculated parameters such as event frequency, lifetime, span, and more.

#### Principal component analysis

Deviations from the rolling mean value for each cell at each frame of time were collected for all the experimental conditions from the 3T3 cell experiments. These were compiled and used to calculate the shape modes using principal component analysis. The shape modes were then used to calculate scalar weights fits based on the cell outline radii.

