## Supplemental Figures for "Comparative profiling of cellular gait on adhesive micropatterns defines statistical patterns of activity that underlie native and cancerous cell dynamics"

Contains:

Supplemental Figures S1-S4 and Legends.

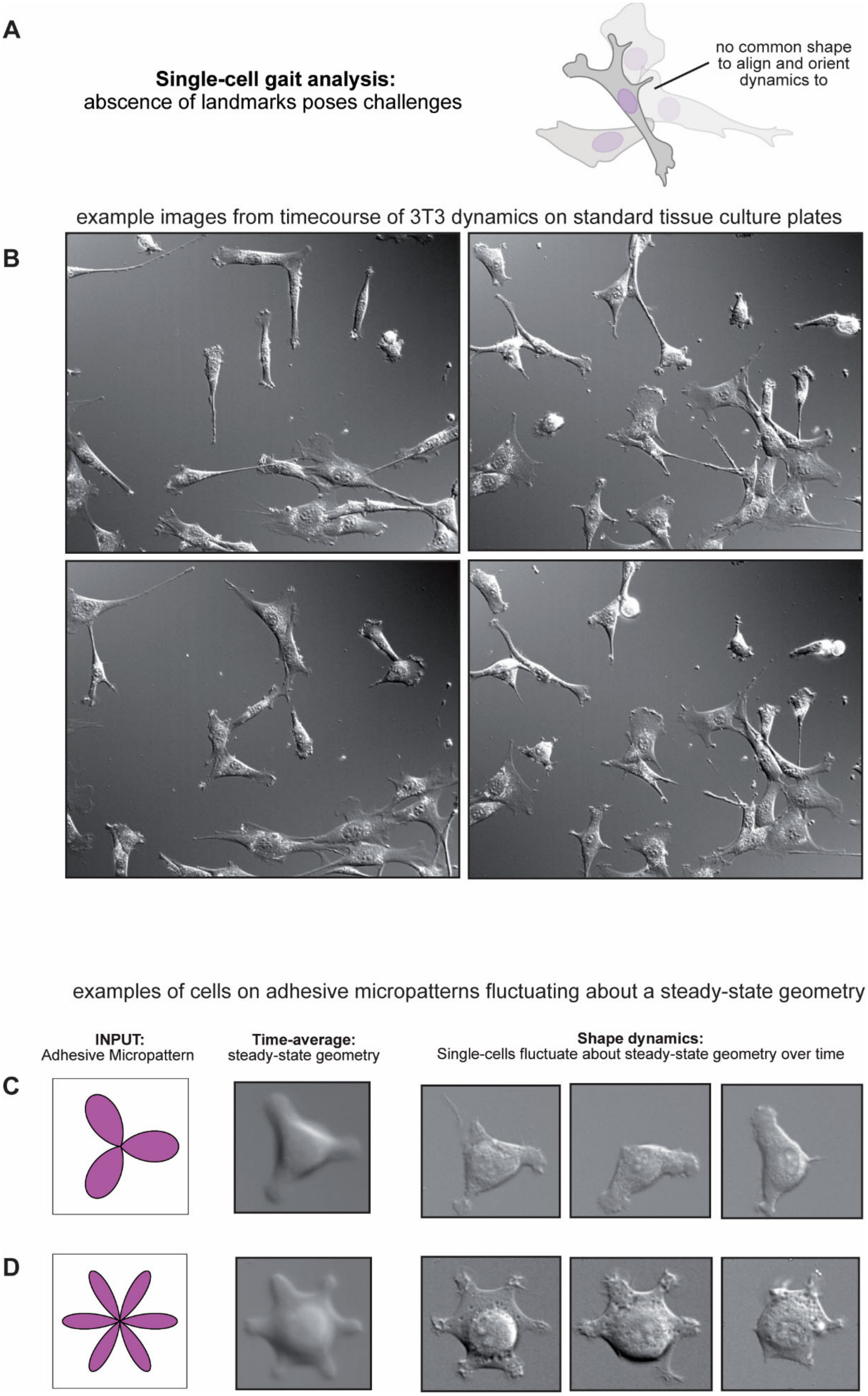
A
Single-cell gait analysis:
abscence of landmarks poses challenges
no common shape to align and orient dynamics to
example images from timecourse of 3T3 dynamics on standard tissue culture plates
B
examples of cells on adhesive micropatterns fluctuating about a steady-state geometry
INPUT:
Adhesive Micropattern
Time-average:
steady-state geometry
Shape dynamics:
Single-cells fluctuate about steady-state geometry over time
C
D

**Figure S1. Additional data: cells grown on adhesive micropattern islands undergo dynamic shape fluctuations about an average cell shape.**

(a) Schematic for how statistical gait analysis is difficult to apply to metazoan cells owing to a lack of landmarks and stereotyped geometry.

(b) Snapshots of 3T3 cells from an overnight imaging experiment. A wide array of shapes and morphologies are observed, making it difficult to quantify and align the dynamics between cells.

(c) Time-average image produced from an image stack of a 3T3 cell grown on a k3 fibronectin micropattern over a 12 hours period, and representative images from the time-series. The cell geometry is observed to fluctuate substantially from the time-average shape throughout the timecourse.

(d) Time-average image produced from an image stack of a 3T3 cell grown on a k6 fibronectin micropattern over a 12 hours period, and representative images from the time-series. The cell geometry is observed to fluctuate substantially from the time-average shape throughout the timecourse.

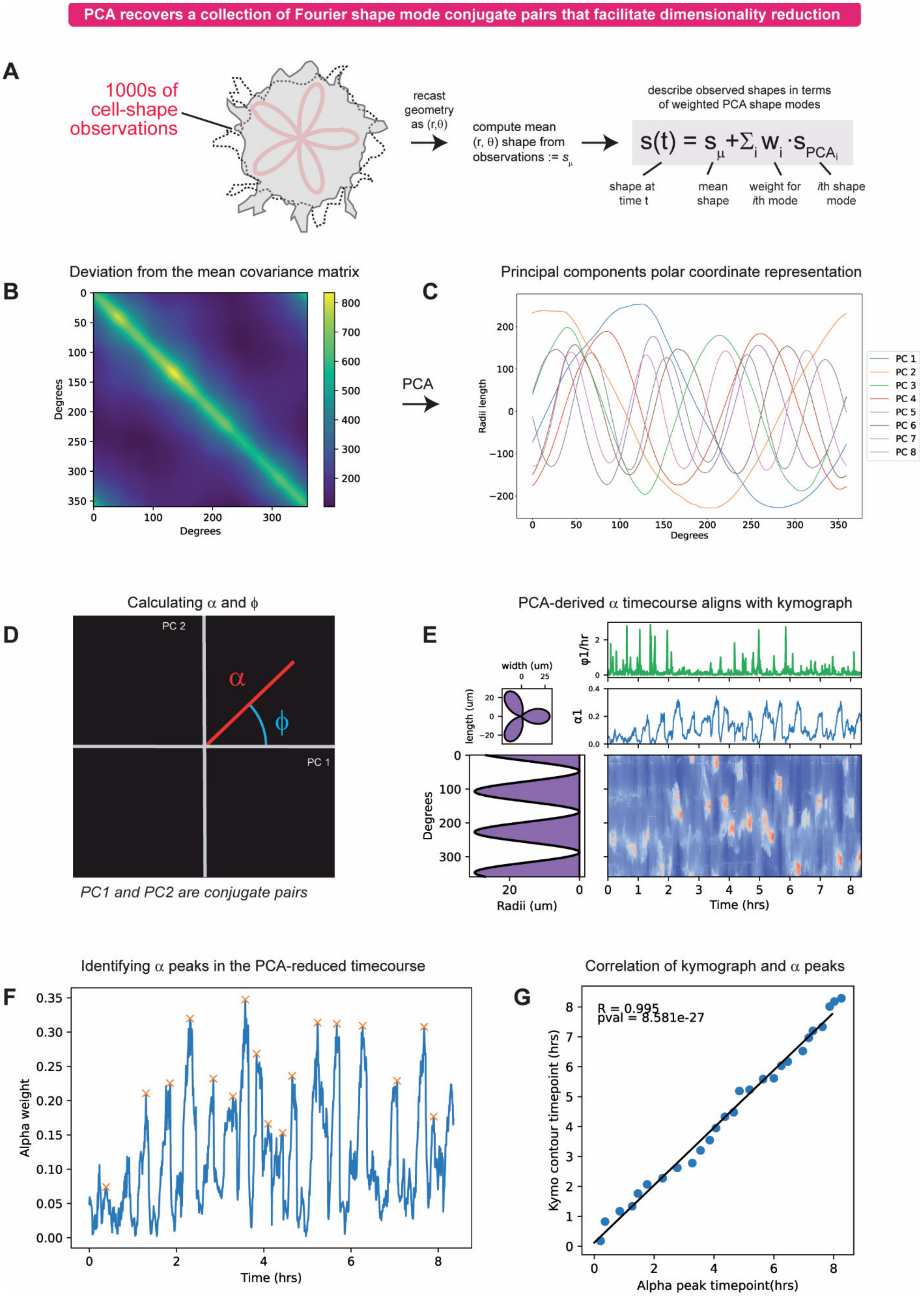

PCA recovers a collection of Fourier shape mode conjugate pairs that facilitate dimensionality reduction
A
1000s of cell-shape observations
recast geometry as (r,θ)
compute mean (r, θ) shape from observations := $s_\mu$
describe observed shapes in terms of weighted PCA shape modes
$s(t) = s_\mu + \Sigma_i w_i \cdot s_{PCA_i}$
shape at time t
mean shape
weight for ith mode
ith shape mode
B
Deviation from the mean covariance matrix
Degrees
Degrees
PCA
C
Principal components polar coordinate representation
Radii length
Degrees
PC 1
PC 2
PC 3
PC 4
PC 5
PC 6
PC 7
PC 8
D
Calculating α and ϕ
PC 2
PC 1
α
ϕ
PC1 and PC2 are conjugate pairs
E
PCA-derived α timecourse aligns with kymograph
width (um)
length (um)
φ1/hr
α1
Degrees
Radii (um)
Time (hrs)
F
Identifying α peaks in the PCA-reduced timecourse
Alpha weight
Time (hrs)
G
Correlation of kymograph and α peaks
R = 0.995
pval = 8.581e-27
Kymo contour timepoint (hrs)
Alpha peak timepoint(hrs)

**Figure S2. Additional data: PCA decomposition and shape-mode fitting.**

(a) Schematic for how a cell's morphological shape dynamics will be described as a deviation from a mean shape in terms of the weights of a collection of shape modes.

(b) Covariance matrix derived from all morphological observations collected.

(c) r,$\theta$ representation of the top 8 PCA-derived shape modes. Note that pairs of conjugate shape modes occur suggesting one can treat these pairs as a fundamental shape mode with an associated magnitude $\alpha$ and phase $\phi$.

(d) Schematic for how the magnitude $\alpha$ and phase $\phi$ are computed from a pair of conjugate shape modes. By treating their combination PCA1+PCA2 as a vector, its magnitude and phase are computed geometrically.

(e) Comparison of PCA-reduced transform of a representative cell's morphological signal to its higher-dimensional kymograph. Time courses for the $\alpha$ parameter, and phase $\phi$ are shown superimposed on the same timescale as the morphological signal kymograph. Note that when high shape mode usage occurs (high $\alpha$ parameter) this leads to a stabilization of the phase $\phi$. Note also that the portions of the timecourse with high $\alpha$ parameter appear to correlate with events in the kymograph.

(f) Results of applying automated peak identification to the PCA-reduced a parameter timecourse. Peaks are denoted with an orange x.

(g) Correlation between appearance of PCA-derived peaks in the $\alpha$ parameter timecourse with events in the kymograph.

### Raw statistics of morphology dynamics within a cell line for different geometries of adhesive micropatterned island

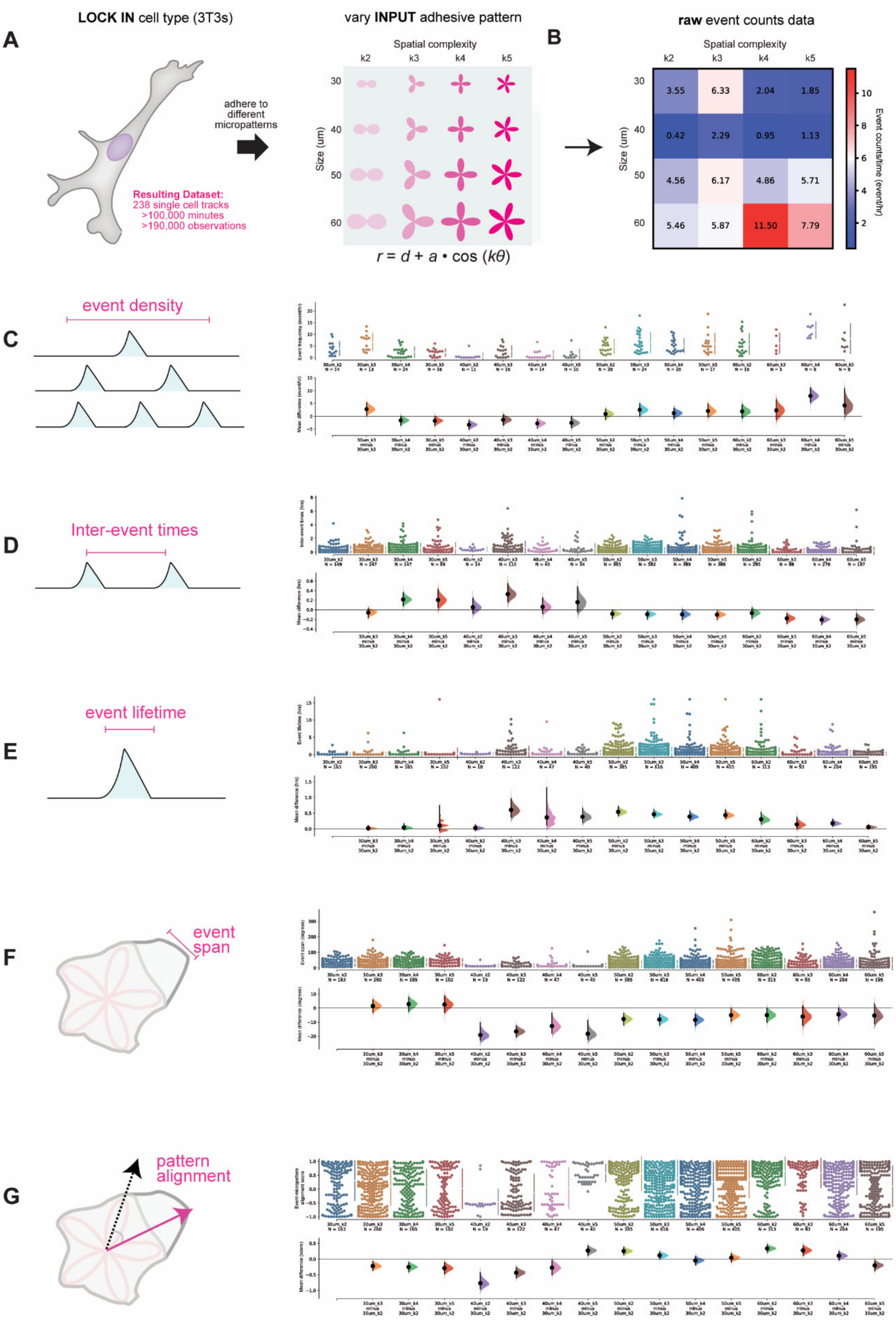

**Figure S3. Additional data: the spatial structure of a micropattern island modulates the statistical features of a cell's dynamic morphology signal.**

(a) Schema depicting experimental design for interrogating effects of micropattern geometry on 3T3 morphology signals. 3T3 cells were imaged on different micropatterns defined by the harmonics of a circle, such that the spatial complexity and radius of the pattern were independently controlled. The resulting single-cell tracks were digitized for analysis using the workflow from Main Text Fig. 2 and Fig. 2S and aggregated by INPUT pattern for statistical comparison between different conditions.

(b) Raw event count density (events per hour) derived from aggregating the data for each of the micropattern geometries tested.

(c) Additional raw data and visualizations of the "event density" distributions from Main Text Fig. 3. Top: raw "event density" distributions across the panel of INPUT patterns tested. Bottom: transformed distributions using the 30 um k2 pattern as the reference for comparison by the "shared control bootstrapping" method.

(d) Additional raw data and visualizations of the "inter-event time" distributions from Main Text Fig. 3. Top: raw "inter-event time" distributions across the panel of INPUT patterns tested. Bottom: transformed distributions using the 30 um k2 pattern as the reference for comparison by the "shared control bootstrapping" method.

(e) Additional raw data and visualizations of the "event lifetime" distributions from Main Text Fig. 3. Top: raw "event lifetime" distributions across the panel of INPUT patterns tested. Bottom: transformed distributions using the 30 um k2 pattern as the reference for comparison by the "shared control bootstrapping" method.

(f) Additional raw data and visualizations of the "event span" distributions from Main Text Fig. 3. Top: raw "event span" distributions across the panel of INPUT patterns tested. Bottom: transformed distributions using the 30 um k2 pattern as the reference for comparison by the "shared control bootstrapping" method.

(g) Additional raw data and visualizations of the "pattern alignment" distributions from Main Text Fig. 3. Top: raw "pattern alignment" score distributions across the panel of INPUT patterns tested. Bottom: transformed distributions using the 30 um k2 pattern as the reference for comparison by the "shared control bootstrapping" method.

### Raw statistics of morphology dynamics for cancer cell lines grown on a panel of adhesive micropatterned islands

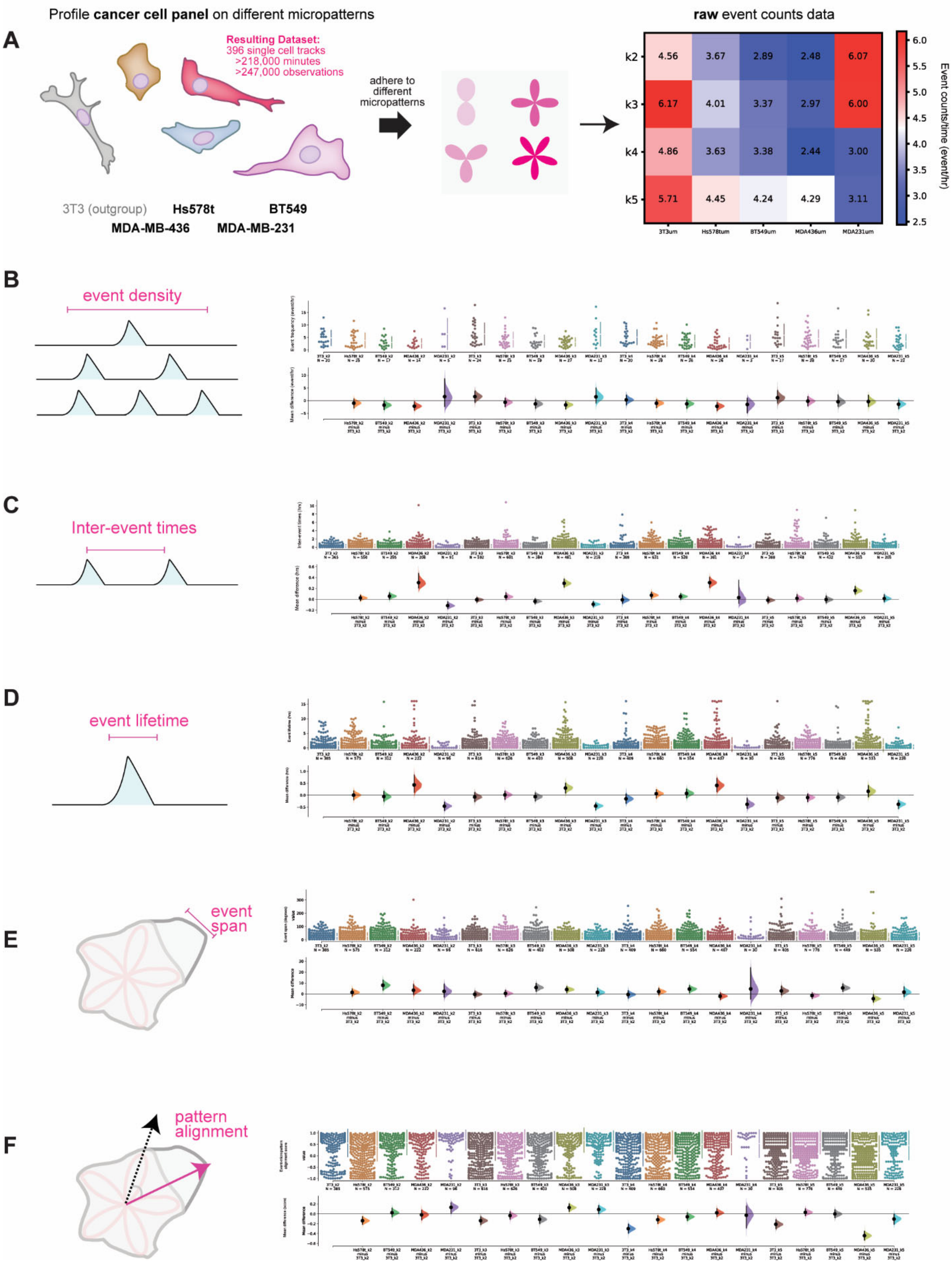

**Figure S3. Additional data: profiling cancer-cell lines on micropatterned islands identifies different statistical axes associated with divergent morphological signals and cell dynamics.**

(a) Left: schema depicting experimental design for building statistical profiles for different cancer cell lines based on the morphology signals they generate when adhered to different micropattern geometries. Cell lines were imaged on 50 um micropatterns across 4 levels of spatial complexity (k2, k3, k4, k5). The resulting single-cell tracks were digitized for analysis using the workflow from Fig. 2 and aggregated by cell-type and INPUT pattern for statistical comparison between different conditions. Right: raw event count density (events per hour) derived from aggregating the data for each of the micropattern geometries and cell lines tested.

(b) Additional raw data and visualizations of the “event density” distributions from Main Text Fig. 4. Top: raw “event density” distributions across the panel of INPUT patterns tested. Bottom: transformed distributions using the 30 um k2 pattern as the reference for comparison by the “shared control bootstrapping” method.

(c) Additional raw data and visualizations of the “inter-event time” distributions from Main Text Fig. 4. Top: raw “inter-event time” distributions across the panel of INPUT patterns tested. Bottom: transformed distributions using the 30 um k2 pattern as the reference for comparison by the “shared control bootstrapping” method.

(d) Additional raw data and visualizations of the “event lifetime” distributions from Main Text Fig. 4. Top: raw “event lifetime” distributions across the panel of INPUT patterns tested. Bottom: transformed distributions using the 30 um k2 pattern as the reference for comparison by the “shared control bootstrapping” method.

(e) Additional raw data and visualizations of the “event span” distributions from Main Text Fig. 4. Top: raw “event span” distributions across the panel of INPUT patterns tested. Bottom: transformed distributions using the 30 um k2 pattern as the reference for comparison by the “shared control bootstrapping” method.

(f) Additional raw data and visualizations of the “pattern alignment” distributions from Main Text Fig. 4. Top: raw “pattern alignment” score distributions across the panel of INPUT patterns tested. Bottom: transformed distributions using the 30 um k2 pattern as the reference for comparison by the “shared control bootstrapping” method.
